# Abnormalities in Proinsulin Processing in Islets from Individuals with Longstanding T1D

**DOI:** 10.1101/542027

**Authors:** Emily K. Sims, Julius Nyalwidhe, Farooq Syed, Henry T. Bahnson, Leena Haataja, Cate Speake, Margaret A. Morris, Raghavendra G. Mirmira, Jerry Nadler, Teresa L. Mastracci, Peter Arvan, Carla J. Greenbaum, Carmella Evans-Molina

## Abstract

Work by our group and others has suggested that elevations in circulating proinsulin relative to C-peptide is associated with development of Type 1 diabetes (T1D). We recently described the persistence of detectable serum proinsulin in a large majority (95.9%) of individuals with longstanding T1D, including individuals with undetectable serum C-peptide. Here we describe analyses performed on human pancreatic sections from the nPOD collection (n=30) and isolated human islets (n=10) to further explore mechanistic etiologies of persistent proinsulin secretion in T1D. Compared to nondiabetic controls, immunostaining among a subset (4/9) of insulin positive T1D donor islets revealed increased numbers of cells with proinsulin-enriched, insulin-poor staining. Laser capture microdissection followed by mass spectrometry revealed reductions in the proinsulin processing enzymes prohormone convertase 1/3 (PC1/3) and carboxypeptidase E (CPE) in T1D donors. Twenty-four hour treatment of human islets with an inflammatory cytokine cocktail reduced mRNA expression of the processing enzymes PC1/3, PC2, and CPE. Taken together, these data provide new mechanistic insight into altered proinsulin processing in long-duration T1D and suggest that reduced β cell prohormone processing is associated with proinflammatory cytokine-induced reductions in proinsulin processing enzyme expression.

## INTRODUCTION

Type 1 diabetes (T1D) is a chronic autoimmune disease that results from immune-mediated destruction of pancreatic β cells, leading to a lifelong dependence on exogenous insulin therapy (1). Classic models of type 1 diabetes pathogenesis have suggested near complete destruction of pancreatic β cells by the time of diagnosis (1). However, recent data from clinical cohorts of individuals with long-duration disease has shown that a significant proportion of individuals with T1D > 3 years duration have detectable C-peptide, and that C-peptide secretion is meal-responsive (2-5). In parallel, histologic analysis of pancreata from organ donors with diabetes has demonstrated the presence of insulin containing islets many years after diagnosis (6, 7). Together, these findings have challenged the notion that all β cells are destroyed in diabetes, while also raising a number of questions regarding the molecular and functional phenotype of β cells that persist in long-duration disease.

The hallmark of a healthy β cell is efficient and robust processing of preproinsulin into mature insulin protein, which is secreted in response to nutrient stimulation. Under normal conditions, preproinsulin is converted to proinsulin following cleavage of its signal peptide by signal peptidases within the lumen of the endoplasmic reticulum (ER) (8). Proinsulin disulfide bond formation and terminal protein folding occur in the ER and Golgi apparatus, and intact proinsulin is eventually cleaved into mature insulin and C-peptide by the enzymes prohormone convertase 1/3 (PC1/3), PC2, and carboxypeptidase E (CPE) in secretory granules (8). In response to inflammatory, oxidative, and endoplasmic reticulum stress in diabetes, protein processing capacity within the secretory compartment of the β cell can become overwhelmed. This leads to the accumulation of inadequately processed proinsulin (9) that can be detected noninvasively in the circulation by measurement of the ratio of proinsulin relative to circulating mature insulin or C-peptide, or the PI:C ratio (10).

We recently described the persistence of detectable serum proinsulin in a majority (95.9%) of individuals with longstanding T1D (≥3 years) who were followed as part of the T1D Exchange Registry (11). Remarkably, this finding extended to individuals who had undetectable serum C-peptide; specifically, 89.9% of individuals without detectable stimulated serum C-peptide had readily detectable serum proinsulin (11). Moreover, even in those with detectable C-peptide, PI:C ratios were significantly increased compared to controls, and higher fasting PI:C ratios were found in individuals with the worst stimulated beta cell function (11). Intriguingly, serum proinsulin levels remained steady over 4 years of follow-up, despite continued loss of C-peptide (11). These findings suggested that in long-duration type 1 diabetes, the ability to secrete proinsulin persists, even in individuals who no longer secrete detectable levels of serum C-peptide. However, the mechanistic etiologies behind these findings have not been elucidated.

In the present study, we analyzed pancreatic sections from donors with established T1D and performed *in vitro* analyses of cytokine treated human islets. Immunostaining analysis of pancreatic sections from T1D donors confirmed discrepancies in proinsulin and C-peptide localization at the level of the islet in some donors with longstanding T1D, while mass spectroscopy analysis of laser-captured β cells from organ donors with diabetes suggested that defects in hormone processing may arise from impaired expression of proinsulin processing enzymes. Consistent with this, cytokine treatment of human islets was associated with reductions in mRNA levels encoding proinsulin processing enzyme. Taken together, these data highlight a prominent role for defective prohormone processing in T1D and provide novel insight into the molecular phenotype of the β cell in longstanding disease.

## METHODS

Human islet sections were obtained through the Network for Pancreatic Organ Donors with Diabetes (nPOD) (12). Sections for immunostaining were obtained from 7 nondiabetic controls and 17 donors with type 1 diabetes. Sections from donors with type 1 diabetes were selected to include a range of diabetes durations, individuals with and without detectable random C-peptide, and documented presence or absence of insulin positive islets. Three of the donors with type 1 diabetes had detectable random serum C-peptide and 13 were classified by nPOD as C-peptide negative (random serum C-peptide <0.017nmol/L via TOSOH immunossay) (12). One donor with type 1 diabetes did not have an available serum C-peptide value. Sections for laser capture microdissection (LCM) were obtained from 3 nondiabetic controls and 3 donors with type 1 diabetes who were known to have positive islet autoantibodies and documented presence of residual insulin positive islets (Case IDs 6234, 6238, 6271, 6212, 6245, and 6247).

Isolated human islets from 10 nondiabetic donors were obtained from the Integrated Islet Distribution Program. Islets from each donor were treated with or without 24 hours of a cytokine cocktail of IL-1β (50 U/ml) and INF-γ (1000 U/ml) (R&D Systems, USA). Total RNA was recovered using RNeasy mini kits (Qiagen, USA), reverse transcribed, and subjected to qRT-PCR using SensiFAST™SYBR Lo-ROX kit (Bioline, USA). Data were analyzed in triplicates, normalized to Glyceraldehyde-3-Phosphate Dehydrogenase GAPDH, and presented as fold change in expression relative to untreated islets from the same individual using the 2^-ΔΔ*CT* method. Primer sequences used were as follows.

### Immunostaining

Tissue sections were deparaffinized through graded xylene and ethanol and permeabilized in PBS containing 0.1% triton-X 100 (FisherScientific). Sections were co-stained with primary antibodies against insulin (guinea pig; 1:500 and 1:1000; Millipore) and proinsulin (mouse; 1:50 and 1:200; Developmental Studies Hybridoma Bank; detects the B-C junction of human proinsulin) as previously described and validated (13, 14). Antii-guinea pig Ig Alexa 647 and anti-mouse Ig Alexa 488 secondary antibodies were used (1:400; Jackson ImmunoResearch). Images were acquired using a LSM 700 confocal microscope (Zeiss).

### Mass spectrometry Analysis of Laser Capture Microdissection (LCM) Islets

As previously described, high resolution high mass accuracy label free quantitative mass spectrometry analysis was applied to islets isolated by laser capture microdissection from nPOD pancreatic sections from 3 donors with type 1 diabetes, and 3 nondiabetic controls (15). To select insulin positive islets for analysis, islets in unstained tissue sections were identified based on intrinsic autofluorescence, which has been shown to correlate with insulin staining (15).

Serial 10-µm-thick tissue sections were prepared from nPOD pancreas blocks and attached onto polyethylene naphthalate (PEN) membrane slides. The tissue sections were dehydrated with graded ethanol solutions before isolating and collecting islets on cups using an ArcturusXT™ LCM instrument (Thermo Fisher Scientific), which is equipped with dual ultraviolet (UV) and infrared (IR) lasers. The IR laser captures cells of interest, while the UV laser microdissects cells of interest and prevents any significant contamination of captured material with adjacent acinar tissue. Islets in unstained tissue sections were identified by their unique and specific intrinsic florescence behavior, which correlates with insulin staining, and was detected using triple filter upon UV illumination of the specimen (16, 17). Protein extraction was performed using the Liquid Tissue MS Protein Prep Kit (Expression Pathology, Rockville, MD) according to the manufacturer’s protocol. Briefly, the LCM cap films containing approximately 3 x10^4^ cell equivalents (estimates based on the thickness and area of the captured islet tissue) were transferred into 20 µl of liquid tissue buffer in a 1.5 ml low protein binding reaction tube and centrifuged at 10,000 × g for 2 minutes to pellet the film. The islet proteins were extracted by heating the mixture at 95°C for 90 minutes with intermittent mixing at 20 minute intervals. After 90 minutes, the samples were centrifuged at 10,000 × g for 1 minute before cooling in ice for 2 min. The equivalent of 1:50 trypsin was added to the extracted protein and incubated at 37°C for 18 hours to generate tryptic peptides. After the trypsin digestion step, an aliquot of the generated peptides was used in a Micro BCA assay to determine the peptide concentrations. The remaining peptides were reduced (10mM DDT) and alkylated (35mM iodoacetamide) and stored at −80°C prior to use for mass spectrometry analysis. LC-MS analysis of digested samples was conducted as previously described (18). Tryptic peptides were solubilized in normalized volumes of 0.1% formic acid (FA)/H2O and the concentration of the digested peptides were determined a Micro BCA assay. The final concentration of the samples was adjusted to 0.5µg/µl using the same buffer. Identical concentrations (2µg) of the peptides were analyzed on a Q-Exactive Orbitrap mass spectrometer coupled to an Easy NanoLC-1000 system (Thermo Fisher Scientific). Each sample was analyzed in triplicate. Protein identification and label-free quantification (LFQ) were performed with MaxQuant software package (18, 19). Analysis of MS spectra was performed using the following parameters: acetylation of the protein N-terminus and oxidation of methionine as variable modifications and carbamidomethylation of cysteine as fixed modification. UniProt-SwissProt human canonical database (version 2016, canonical proteome; 20 198 identifiers) was selected as FASTA file. Seven amino acids were selected as minimum peptide length. Mass accuracy thresholds for the analysis were as follows: 15ppm for MS and 0.8 Da for MS/MS. Match between runs option was kept as default (match time window: 0.7 min; alignment time window: 4 min). LFQ was enabled and LFQ minimum ratio count was set to 1. Remaining options were kept as default. Only unique and razor peptides were used for identification and quantitation. Mascot was also used for database searches and identification (Matrix Science). Tandem mass MS2 spectra identifying and validating the peptides used for the validation of quantitation of each protein are displayed below in Figure 1.

**Figure 1.**
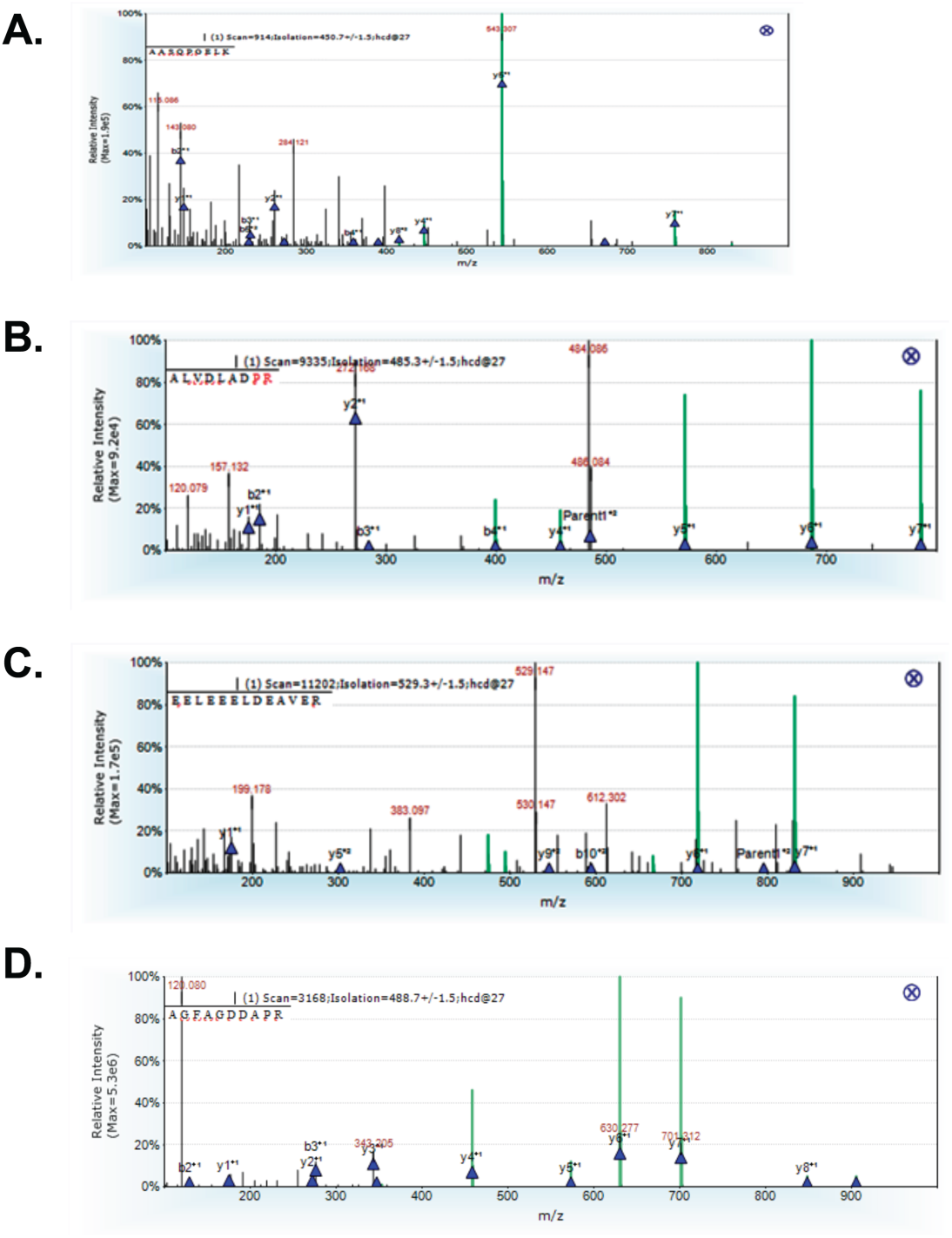
Tandem mass MS2 spectra identifying and validating the peptides used for the validation of quantitation of each protein in global islet proteomics analysis. **A.** Tandem mass spectra identifying and validating the peptide used for quantitation of carboxypeptidase E (AASQPGELK). The corresponding y and b ions are marked with blue triangles. The Mascot ion score for this peptide was 34 with an expect score of 0.0098. **B.** Tandem mass spectra identifying and validating the peptide used for quantitation of prohormone convertase 1/3 (ALVDLADPR). The corresponding y and b ions are marked with blue triangles. The Mascot ion score for this peptide was 36 with an expect score of 0.0063. **C.** Tandem mass spectra identifying and validating the peptide used for quantitation of prohormone convertase 2 (EELEEELDEAVER). The corresponding y and b ions are marked with blue triangles. The Mascot ion score for this peptide was 32 with an expect score of 0.012. **D.** Tandem mass spectra identifying and validating the peptide used for quantitation of actin (AGFAGDDAPR). The corresponding y and b ions are marked with blue triangles. The Mascot ion score for this peptide was 70 with an expect score of 9.17E-7.

### Statistics

Analyses of cytokine treated islet data were performed using GraphPad Prism 7.0. A student’s t-test was used to compare expression of proinsulin processing enzymes between control and cytokine treated islets.

For proteomic analyses, Perseus (version 1.5.2.6) was used to perform statistical analysis of LFQ proteomic data after Log_2_ transformation, data imputation and filtration. Two-sample t-tests were used together with permutation-based false discovery rate calculations for comparative quantitative analysis of protein expression in the two sample cohorts (T-test, FDR= 0.05, *p* < 0.05). This analysis takes into consideration all the peptides that are identified from the protein. Pinnacle, a quantitative proteomic analysis software (Optys Tech Corporation) was used for visualization and validation of the LFQ data using the areas under the curve from extracted ion chromatograms of precursor ions of unique peptides from targeted proteins. Pinnacle utilizes extracted ion chromatograms generated from the mass spectrometry MS1 data feature (peptides) and their corresponding MS2 fragmentation data to confirm the identity of the signal for each replicate sample. Peptides with the best ionization efficiencies, peak shape and isotopic fidelity were used for comparative analysis. Actin was used as an internal control. Unpaired t-tests were used to compare mean area under the curve values for each subject group. For all analyses, two-tailed p values of ≤0.05 were considered significant.

## RESULTS

To explore whether abnormalities in proinsulin processing were present in pancreatic tissue sections from 17 donors with type 1 diabetes from the nPOD collection, immunostaining was performed for insulin and proinsulin in pancreatic sections from nondiabetic control donors and donors characterized as having clinical type 1 diabetes (duration 1.5-28 years) with a wide range of demographic and clinical characteristics (Table 2). In control donors, the majority of β cells exhibited both proinsulin and insulin staining with limited overlap observed between regions of proinsulin and insulin staining. Within our cohort of donors with T1D, 47% (n=8) lacked any detectable insulin and proinsulin staining. Among sections from donors with T1D with insulin positive cells, many sections exhibited scattered insulin positive cells or islets lacking central insulin positive cells. Although 55% (n=5) of these insulin positive sections exhibited a staining pattern where the majority of insulin+ β-cells also stained positive for proinsulin, in 4 donors (Table 2, Figure 2; Case IDs 6040, 6069, 6243, and 6211), a different phenotype was seen where multiple proinsulin-enriched but insulin poor β-cells were observed.

**Table 1.**
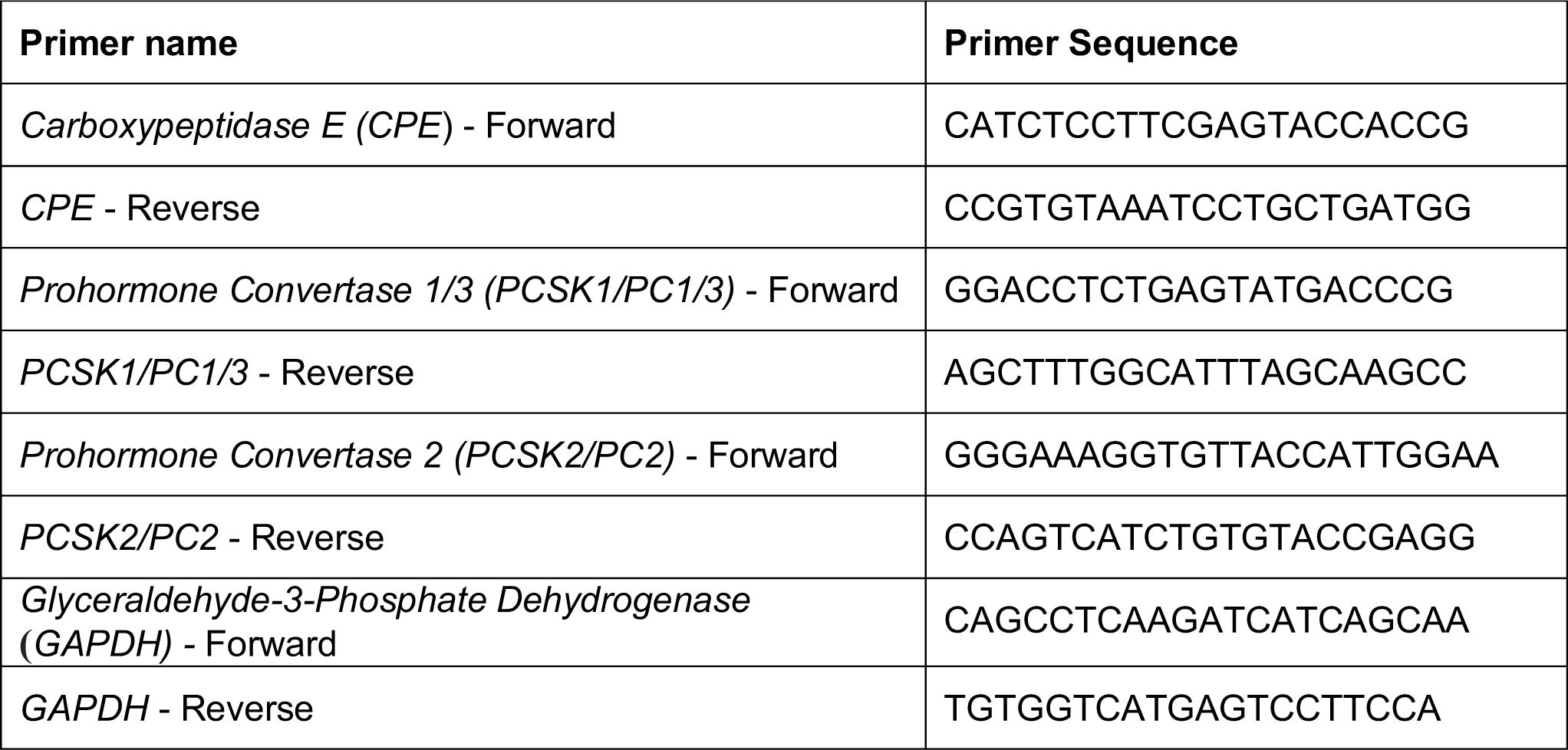
Primer Sequences.

**Table 2.** Characteristics of nPOD Donor Staining.

| **Case ID** | **Age** | **T1D Duration (years)** | **Sex** | **BMI** | **Positive aAbs** | **Detectable Fasting Serum C-peptide** | **Insulin + Cells Present** | **Proinsulin (PI) Staining Phenotype** |
| --- | --- | --- | --- | --- | --- | --- | --- | --- |
| **6015** | 39 | n/a (Control) | F | 32.2 | n/a | n/a | + | Multiple costaining cells |
| **6029** | 24 | n/a (Control) | F | 22.6 | n/a | n/a | + | Multiple costaining cells |
| **6034** | 32 | n/a (Control) | F | 25.2 | n/a | n/a | + | Multiple costaining cells |
| **6055** | 27 | n/a (Control) | M | 22.7 | n/a | n/a | + | Multiple costaining cells |
| **6104** | 41 | n/a (Control) | M | 20.5 | n/a | n/a | + | Multiple costaining cells |
| **6179** | 20 | n/a (Control) | F | 21.8 | n/a | n/a | + | Multiple costaining cells |
| **6229** | 31 | n/a (Control) | F | 26.9 | n/a | n/a | + | Multiple costaining cells |
| **6038** | 37.2 | 20 | F | 30.9 | - | + | + | Multiple costaining cells |
| **6041** | 26.3 | 23 | M | 28.4 | - | - | - | None |
| **6054** | 35.1 | 30 | F | 30.4 | mIAA | - | - | None |
| **6077** | 32.9 | 19 | F | 22 | mIAA | - | - | None |
| **6141** | 36.7 | 28 | M | 26 | mIAA, IA-2, GAD, ZnT8 | - | - | None |
| **6143** | 32.6 | 7 | F | 26.1 | mIAA, IA-2 | - | - | None |
| **6173** | 44.1 | 15 | M | 23.9 | - | - | - | None |
| **6180** | 27.1 | 11 | M | 25.9 | mIAA, IA-2, GAD, ZnT8 | - | - | None |
| **6208** | 32.6 | 16 | F | 23.4 | - | - | - | None |
| **6040** | 50 | 20 | F | 31.6 | mIAA | - | + | Pseudoatrophic islets with rare insulin+ cells which were mostly PI-enriched, insulin poor cells |
| **6069** | 22.9 | 7 | M | 28.8 | Unknown | Unknown | + | Mixed beta cell phenotype with some islets displaying mostly costaining cells and some islets with numerous PI-enriched, insulin poor cells |
| **6070** | 22.6 | 7 | F | 21.6 | mIAA, IA-2 | - | + | Multiple costaining cells |
| **6121** | 33.9 | 4 | F | 18 | - | + | + | Multiple costaining cells |
| **6211** | 24 | 4 | F | 24.4 | miAA, IA-2,<br>GAD, ZnT8 | - | + | Occasional PI-enriched, insulin-poor cells |
| **6224** | 21 | 1.5 | F | 22.8 | - | - | + | Multiple costaining cells |
| **6237** | 18 | 12 | F | 26 | miAA, GAD | - | + | Rare isolated insulin+ cells, all costain |
| **6243** | 13 | 5 | M | 21.3 | miAA | + | + | Clusters of PI-enriched insulin-poor cells |

nPOD- Network for Pancreatic Organ Donors with Diabetes; T1D- Type 1 Diabetes Mellitus; BMI- Body Mass Index; aAbs-
autoantibodies; proinsulin-PI; miAA- microinsulin autoantibody; IA-2- islet antigen 2 antibody; GAD- glutamic acid decarboxylase
antibody; ZnT8- zinc transporter 8 antibody

**Figure 2:**
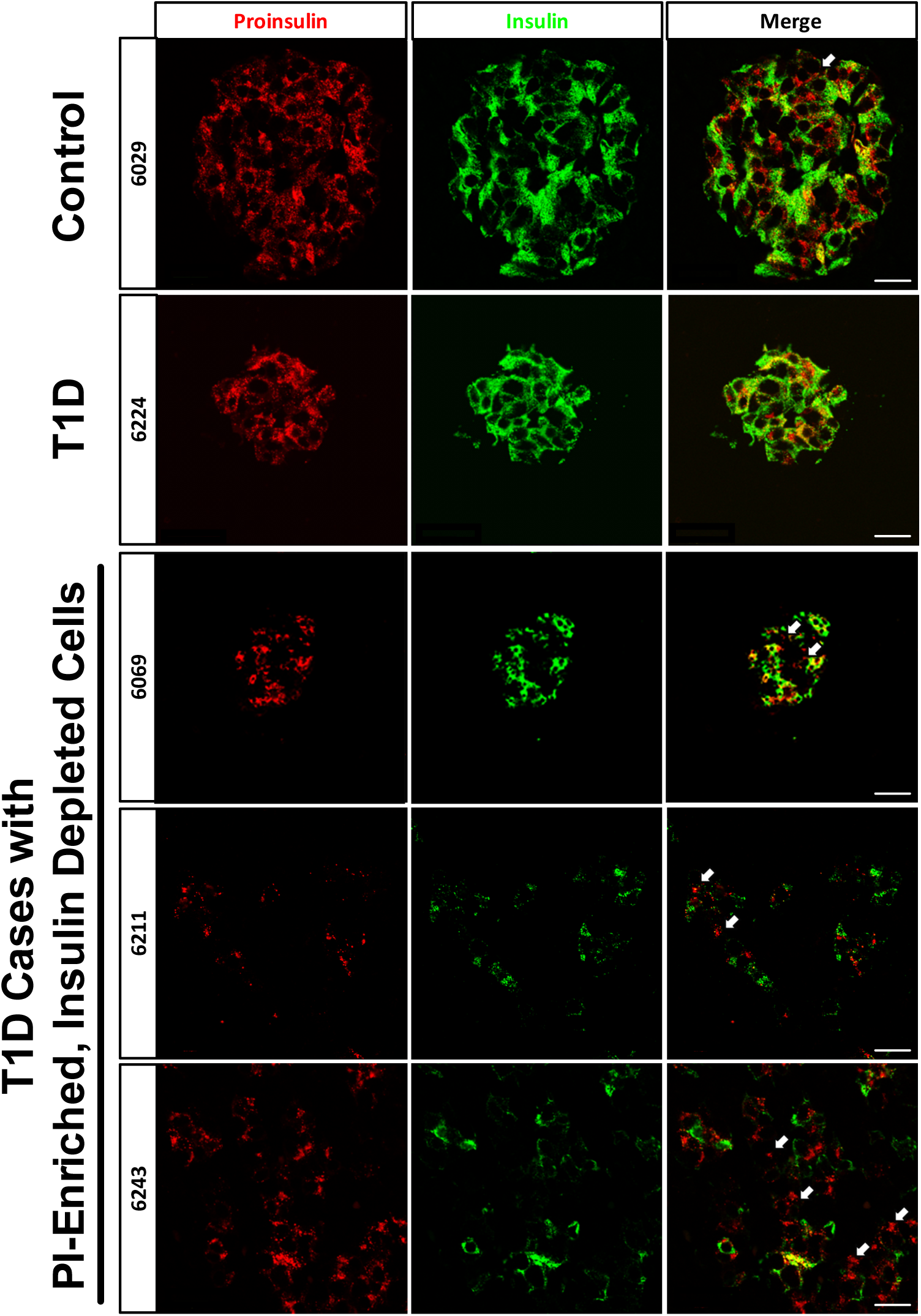
Examples of Proinsulin (PI) and Insulin Immunostaining Patterns in Type 1 Diabetes (T1D) Donor Islets. Immunostaining of proinsulin (green) and insulin (red) was performed on pancreata from non-diabetic control donors and 16 T1D donors. Staining from 3 donors exhibiting multiple PI-enriched, and insulin depleted cells are shown (indicated by white arrows; case IDs 6069, 6243, 6211). These cells were rare in nondiabetic control donors. Scale bars represent 200 µm.

To explore potential mechanisms of impaired prohormone processing, LCM followed by quantitative mass spectrometry analysis of proinsulin processing enzymes was performed on pancreatic sections from nPOD donors with type 1 diabetes compared to nondiabetic controls. Consistent with a reduction in proinsulin processing, LFQ analysis demonstrated significant reductions in CPE (−3.20 fold change, p<0.001), as well as PC1/3 levels (−1.66 fold change, p<0.001) in islets from donors with type 1 diabetes compared to controls. In contrast, there were no significant differences in expression levels of PC2 (−0.442 fold change; p=0.368) or the control protein, actin (−0.007 fold change; p=0.939), between the two groups. The area under the curve values from extracted ion chromatograms of the MS1 features of representative peptides from the target proteins are quantified in Figure 3A-E. Peptide scores for each of the four peptides were beyond the significance threshold in Mascot searches. Similar to the LFQ analysis, comparison of mean area under the curve values for control donors compared to donors with type 1 diabetes revealed a clear trend towards a reduction in CPE (p=0.0514) and a significant reduction in PC1/3 (p=0.0046) (Figure 3B-C), but no differences in PC2 or actin expression (Figure 3D-E).

**Figure 3.**
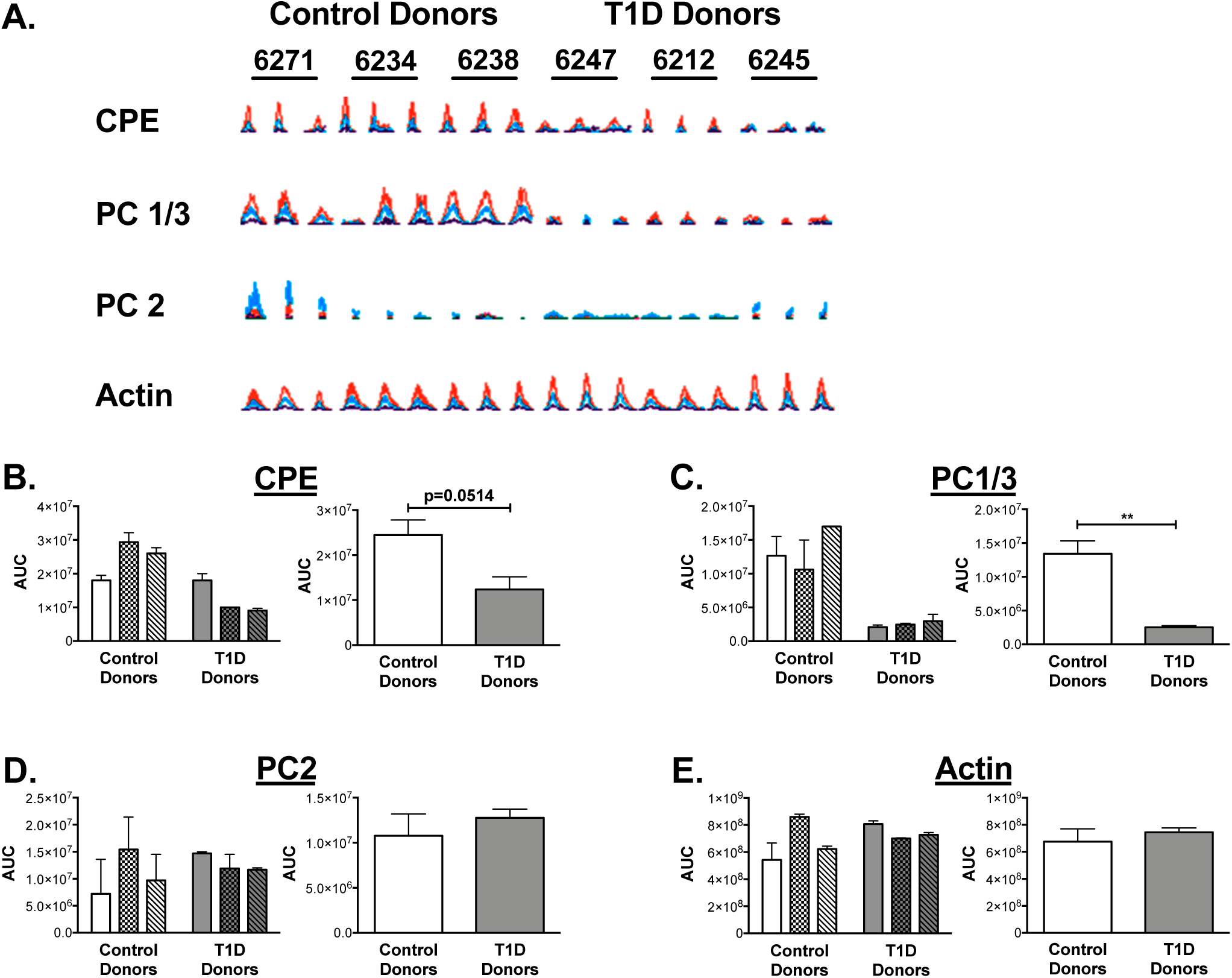
Expression of CPE, PC1/3 and PC2 in LCM islets from nondiabetic controls and donors with type 1 diabetes (T1D). **A.** Extracted ion chromatograms of peptides from nondiabetic individuals (n=3) compared to T1D donors (n=3) with sequences AASQPGELK, ALVDLADPR, EELEEELDEAVER and AGFAGDDAPR from CPE, PC1/3, PC2, and actin, respectively. The Mascot identification ion and expect scores were significant for each of the four peptides. Expression of the individual peptides in each nPOD case (run as triplicates) are shown. nPOD case IDs are indicated above each set of triplicates. **B-E:** Quantitation of area under the curve (AUC) for chromatograms from each peptide is shown for each donor on the left, as well as in aggregate form on the right, with means and standard errors indicated. Aggregate means of control donors were compared to donors with type 1 diabetes using an unpaired t-test. (n=3); **p<0.01 CPE-carboxypeptidase E; PC1/3-prohormone convertase 1/3; PC2-prohormone convertase 2.

To test whether reductions in islet proinsulin processing enzyme expression may be associated with islet inflammatory stress, we assayed expression of CPE, PC1/3, and PC2 in human islets from ten donors that were treated with 24 hours of a cytokine cocktail (50 U/ml IL-1β and 1000 U/ml INF-γ) for 24 hours. As shown in Figure 4, cytokine treatment was associated with significant reductions in mRNA expression for each of the processing enzymes.

**Figure 4.**
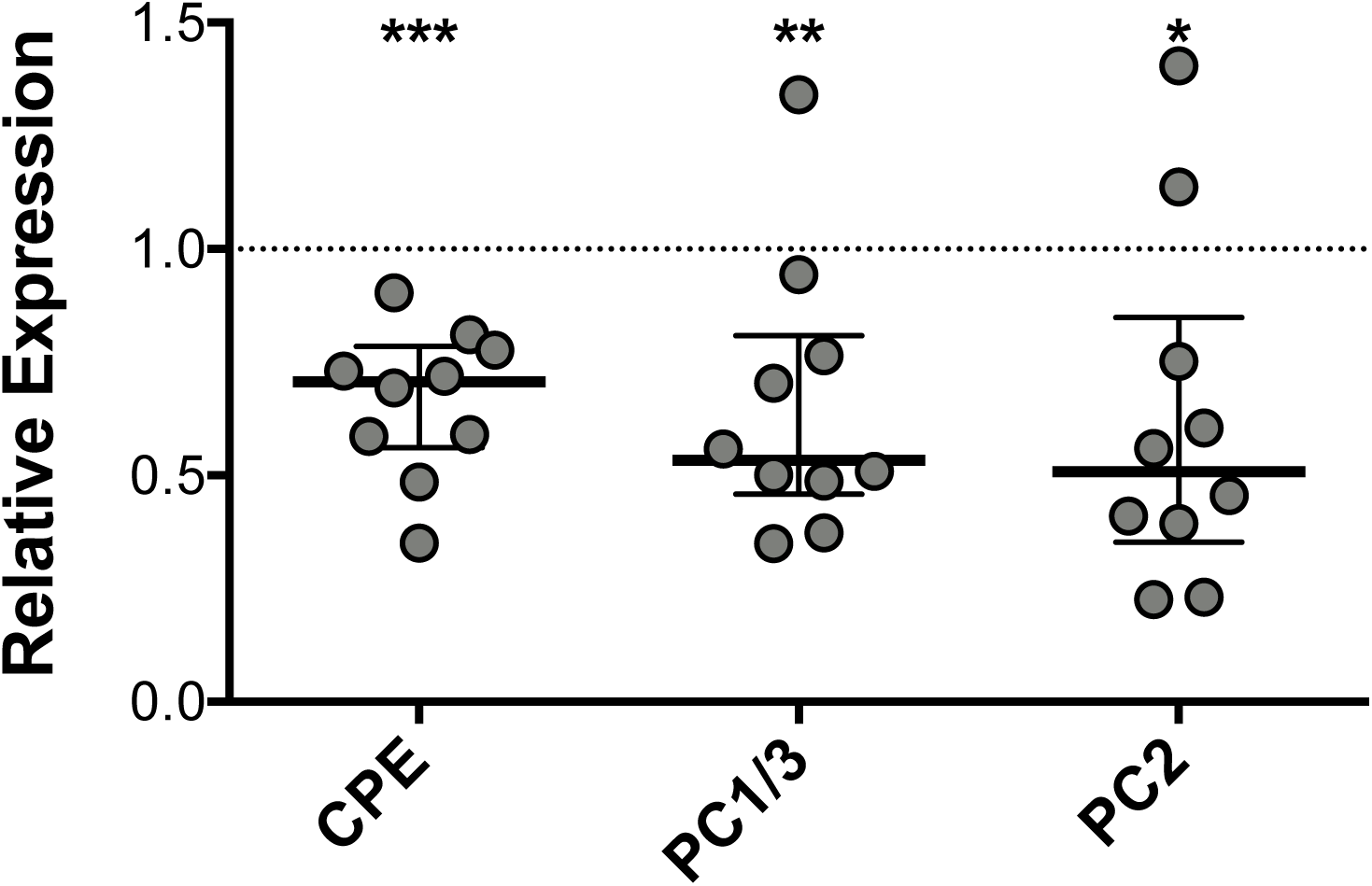
Islet Inflammatory Stress is associated with reductions in proinsulin processing enzyme expression. qRT-PCR for *Carboxypeptidase E (CPE), Prohormone Convertase 1/3 (PC1/3),* and *Prohormone Convertase 2 (PC2)* was performed in human islets treated with or without an inflammatory cytokine cocktail consisting of of IL-1β (50 U/ml) and INF-γ (1000 U/ml) for 24 hours. Bars shown represent median +/-interquartile range of mRNA expression relative to untreated islets from the same donor (indicated by the dashed line). N=10; *p<0.05; **p<0.01; ***p<0.001

## DISCUSSION

We recently reported that the ability to initiate preproinsulin production and secrete proinsulin persists in most individuals with longstanding type 1 diabetes, even in those who were functionally C-peptide negative. Moreover, analysis of longitudinal data from >90 individuals with T1D of ≥3 years duration showed that serum proinsulin levels remained fairly stable over four years of follow-up (11). Here, we show for the first time immunostaining data that identifies a discordance between proinsulin production and conversion to mature insulin within islets of a subset of individuals with established type 1 diabetes, as well as proteomic analysis suggesting that this discordance is correlated with a reduction in the protein expression level of PC1/3 and CPE in islets from type donors with 1 diabetes. These data may help to explain at a mechanistic level the deficiency of C-peptide (and insulin) from individuals that are otherwise proinsulin-positive.

The accumulation of incompletely processed proinsulin in longstanding T1D demonstrates a clearly disturbed environment for proinsulin maturation, a process normally regulated by the processing enzymes, PC1/3, 2, and CPE (8). A previous proteomic analysis of whole pancreas tissue lysates from diabetic donors reported a 10-fold decrease in CPE expression compared to nondiabetic donors (20). Our LCM analysis extends this observation by showing that reductions in the expression of proinsulin processing enzymes are specific to islets in individuals with T1D. Although loss of β-cell mass itself could lead to an absolute reduction in proinsulin processing enzymes, in our study LCM analysis was only performed on islets that contained insulin. This, in combination with the observation that cytokine stress directly lowers the expression of mRNAs encoding prohormone processing enzymes in human islets, provides an additional (alternative, although not mutually exclusive) explanation for our findings. Prohormone convertase and CPE activities are also involved in proamylin processing, and evidence of altered pro islet amyloid polypeptide processing has also been identified in patients with type 1 diabetes, which lends further credence to our observations (21, 22).

Regarding our in vitro data suggesting that diminished proinsulin processing enzyme expression may be, at least in part, associated with islet inflammatory stress during T1D, we recognize that the etiology of these observations could be multifactorial. For one, prior reports have described increased proinsulin release and decreased PC1/3 and PC2 protein in human islets treated with inflammatory cytokines (23). Enzyme expression and/or activity could be also impacted by hyperglycemia or other extrinsic factors, as well as cell intrinsic factors such as activation of ER stress, a pathway increasingly implicated in type 1 diabetes pathogenesis (24, 25). Along these lines, reduced expression, translation, and posttranslational processing of CPE and PC1/3 have been previously been reported in *ex vivo* models of ER stress in human islets (26-28). Alternatively, CPE has been identified previously as an islet autoantigen (29), and we cannot exclude that autoimmune targeting of processing enzymes could lead to altered expression and/or activity of these enzymes. Altered prohormone processing could also represent an inherited phenotype that is present at baseline in some subjects who go on to develop type 1 diabetes, potentially exacerbating progression of the disease. Consistent with this model, polymorphisms near the gene locus encoding CPE have been linked to T1D susceptibility (30).

The increasing identification of persistent β cells in long-duration T1D raises a number of questions regarding the source of these cells and how their molecular phenotype intersects with ambient states of immune activation. To date, analysis of persistent β cells in sections from donors with long-duration T1D using confocal microscopy has not provided strong evidence for β cell replication or neogenesis (7). Key unanswered questions are whether these persistent β cells represent de-differentiated β cells and whether the process of de-differentiation may enable β cells to escape immune recognition. In this regard, a subset of β cells with reduced immunogenicity was identified in the non-obese diabetic (NOD) mouse model of T1D. These cells had reduced insulin content and reduced expression of genes associated with β cell identity, whilst genes associated with immune modulation and “stemness” were increased (31). Interestingly, β-cell dedifferentiation due to transcription factor FoxO1 deletion in mouse models has also been linked also to dramatic reductions in the expression of proinsulin processing enzymes (32). The recent observation that many islets in pancreatic sections from individuals with longstanding T1D harbor very low level insulin content in association with identity markers for both β cells and other islet cells also supports the possibility of dedifferentiated islet cells in longstanding T1D (33). Taken together, our results and the above published findings suggest potential associations between altered proinsulin processing, loss of prohormone processing enzyme expression, and β-cell identity (32). However, additional work is needed in human samples to fully elucidate these relationships.

In summary, our findings demonstrate that persistent circulating proinsulin in individuals with longstanding type 1 diabetes could be related to altered proinsulin processing at the level of the islet, which is exacerbated by islet inflammatory stress. These findings present the tantalizing proposition that such residual proinsulin producing cells are “sleeping” and could potentially be amenable to a therapeutic strategy repairing the processing deficit and “waking” the cells. Along these lines, our data suggest that, in longstanding type 1 diabetes, measurement of proinsulin in addition to C-peptide may provide a more sensitive indicator of persistent β-cell mass than measurement of C-peptide alone. Future work is needed to fully characterize the molecular phenotype of these persistent β cells and to delineate whether therapeutics targeting proinsulin processing could increase endogenous insulin production in type 1 diabetes.

## ACKNOWLEDGMENTS

None of the authors have any relevant conflicts of interest to disclose. E.S. and C.E.M serve as guarantors of this work.

A portion of this research was performed with the support of the Network for Pancreatic Organ Donors with Diabetes (nPOD), a collaborative type 1 diabetes research project sponsored by JDRF. Organ Procurement Organizations (OPO) partnering with nPOD to provide research resources are listed at http://www.jdrfnpod.org/for-partners/npod-partners/. We also acknowledge Wojciech Grzesik for technical assistance with proinsulin measurement and LCM capture.

Other Funding Sources: This manuscript was supported by funding from NIDDK K08DK103983 to E.K.S., a Pediatric Endocrine Society Clinical Scholar Award to E.K.S., and NIH grants R01 DK093954 (to C.E-M.) R01 DK48280 (to P.A) and UC4 DK 104166 (to C.E-M. and R.G.M.), VA Merit Award I01BX001733 (to C.E-M.), JDRF Pioneer Award and Strategic Research Agreement (to C.E-M.), JDRF 47-2014-299-Q-R (C.E-M.), and JDRF grant number 2-SRA-2017-498-M-B (to E.K.S).

